# A Mutation in *Tmem135* Causes Progressive Sensorineural Hearing Loss

**DOI:** 10.1101/2024.05.09.593414

**Authors:** Mi-Jung Kim, Shion Simms, Ghazaleh Behnammanesh, Yohei Honkura, Jun Suzuki, Hyo-Jin Park, Marcus Milani, Yukio Katori, Jonathan E Bird, Akihiro Ikeda, Shinichi Someya

## Abstract

Transmembrane protein 135 (TMEM135) is a 52 kDa protein with five predicted transmembrane domains that is highly conserved across species. Previous studies have shown that TMEM135 is involved in mitochondrial dynamics, thermogenesis, and lipid metabolism in multiple tissues; however, its role in the inner ear or the auditory system is unknown. We investigated the function of TMEM135 in hearing using wild-type (WT) and *Tmem135*^FUN025/FUN025^ (*FUN025*) mutant mice on a CBA/CaJ background, a normal-hearing mouse strain. Although *FUN025* mice displayed normal auditory brainstem response (ABR) at 1 month, we observed significantly elevated ABR thresholds at 8, 16, and 64 kHz by 3 months, which progressed to profound hearing loss by 12 months. Consistent with our auditory testing, 13-month-old *FUN025* mice exhibited a severe loss of outer hair cells and spiral ganglion neurons in the cochlea. Our results using BaseScope *in situ* hybridization indicate that TMEM135 is expressed in the inner hair cells, outer hair cells, and supporting cells. Together, these results demonstrate that the *FUN025* mutation in *Tmem135* causes progressive sensorineural hearing loss, and suggest that TMEM135 is crucial for maintaining key cochlear cell types and normal sensory function in the aging cochlea.

## 1. Introduction

Transmembrane proteins (TMEMs) are involved in diverse biological processes such as energy production, signal transduction, cell-cell interaction, cell-cell communication, and synaptic transmission (Schmit and Michiels, 2018; Marx et al., 2020). TMEM135, also known as PMP52, is a 52 kDa protein with five predicted transmembrane domains that is highly conserved across species (Exil et al., 2010; Lee et al., 2016; Beasley et al., 2021). TMEM135 is implicated in various human diseases, including visual loss, osteoporosis, obesity, nonalcoholic fatty liver disease, and hypertrophic cardiomyopathy (Beasley et al., 2021). TMEM135 is thought to have a role in lipid metabolism and homeostasis: in *C. elegans*, deletion of *tmem135* causes a reduction in fat stores and decreased longevity, while *C*. *elegans* transgenic animals overexpressing TMEM-135 exhibit increased longevity upon exposure to cold stress (Exil et al., 2010). In mice, a point mutation in *Tmem135* impairs lipid metabolism in the eye (Landowski et al., 2022) and regulates the docosahexaenoic acid (DHA) metabolism with the liver, retina, heart, and plasma (Landowski et al., 2023). In human cell lines, TMEM135 is a potential candidate for regulating intracellular cholesterol transport through lysosome-peroxisome membrane contacts (Chu et al., 2015), and regulates primary ciliogenesis through the modulation of intracellular cholesterol distribution (Maharjan et al., 2020).

There is growing evidence that TMEM135 is involved in mitochondrial dynamics. A point mutation in *Tmem135* results in an imbalance of mitochondrial fission and fusion and leads to age-dependent retinal pathologies in mouse (Lee et al., 2016). A subsequent study has shown that an adipose-tissue specific deletion of TMEM135 blocks mitochondrial fission, impairs thermogenesis, and increases diet-induced obesity and insulin resistance in mice, while TMEM135 overexpression promotes mitochondrial fission, counteracts obesity and insulin resistance, and rescues thermogenesis in peroxisome-deficient mice (Hu et al., 2023). The effects of TMEM135 appear conserved in other tissues, with TMEM135 deficiency impairing mitochondrial fission and disrupting crucial mitochondrial energy metabolism during osteogenesis in bone marrow mesenchymal stem cells, and *Tmem135* knockout mice displaying an osteoporotic phenotype, characterized by reduced osteogenesis and increased adipogenesis (Liu et al., 2024).

In this present study, we explore the potential role of Tmem135 in the auditory system. Previous work has shown that *Tmem135* is expressed in inner hair cells, outer hair cells, supporting cells, and stria vascularis (Liu et al., 2018; Kolla et al., 2020), however its function in the inner ear is unknown. We have explored this question using the *Tmem135* (*FUN025*) mutant mouse and show that this loss-of-function allele causes progressive sensorineural hearing loss. Our data demonstrate that TMEM135 is necessary for maintaining key cochlear cell types and more broadly for normal sensory function in the aging cochlea.

## 2. Methods

### 2.1. Animals

Generation and characterization of *Tmem135*^FUN025/FUN025^ mice have been described (Lee et al., 2016). *Tmem135*^FUN025/+^ mice on the CBA/CaJ background were a gift from Dr. Akihiro Ikeda (University of Wisconsin-Madison). CBA/CaJ mice were purchased from the Jackson Laboratory (Jackson Laboratory, 000654). The *Tmem135*^FUN025/FUN025^ mutant mice were originally generated on a C57BL/6J background (Lee et al., 2016). The C57BL/6J-*Tmem135*^FUN025/+^ mice were backcrossed onto the CBA/CaJ mice for four generations and CBA/CaJ-*Tmem135*^FUN025/+^ males were mated with CBA/CaJ-*Tmem135*^FUN025/+^ females to obtain CBA/CaJ-*Tmem135*^+/+^, CBA/CaJ-*Tmem135*^FUN025/+^, and CBA/CaJ-*Tmem135*^FUN025/FUN025^ mice. Male and female CBA/CaJ-*Tmem135*^+/+^ (wild-type), CBA/CaJ-*Tmem135*^FUN025/+^ (heterozygous), and CBA/CaJ-*Tmem135*^FUN025/FUN025^ (homozygous or *FUN025*) littermates were used in the current study. All animal experiments were conducted under the protocols approved by the University of Florida Institutional Animal Care and Use Committee.

### 2.2. Genotyping and Backcrossing

*FUN025 genotyping*. Genomic DNA was isolated from mouse tail biopsy. Primer sequences for PCR were as follows (Lee et al., 2016):

Forward 5’-GGTTTTTGGCAGGTGTGTC-3’;

Reverse 5’-TGTGTGCTTGGCAACTTCTC-3’.

These primers were used for amplification of the WT allele (118 bp) and the *FUN025* allele (118 bp). Subsequently, the restriction enzyme Cac8I (New England Biolabs, R0579L) was used to digest the *FUN025* allele specifically to generate two bands (80 and 38 bp) (Fig. 1*a*). Cycling conditions for PCR were as follows: 95 °C for 2 min; 50 cycles of 94 °C for 20 sec, 50 °C for 30 sec, 72 °C for 40 sec; 72 °C for 7 min.

**Figure 1.**
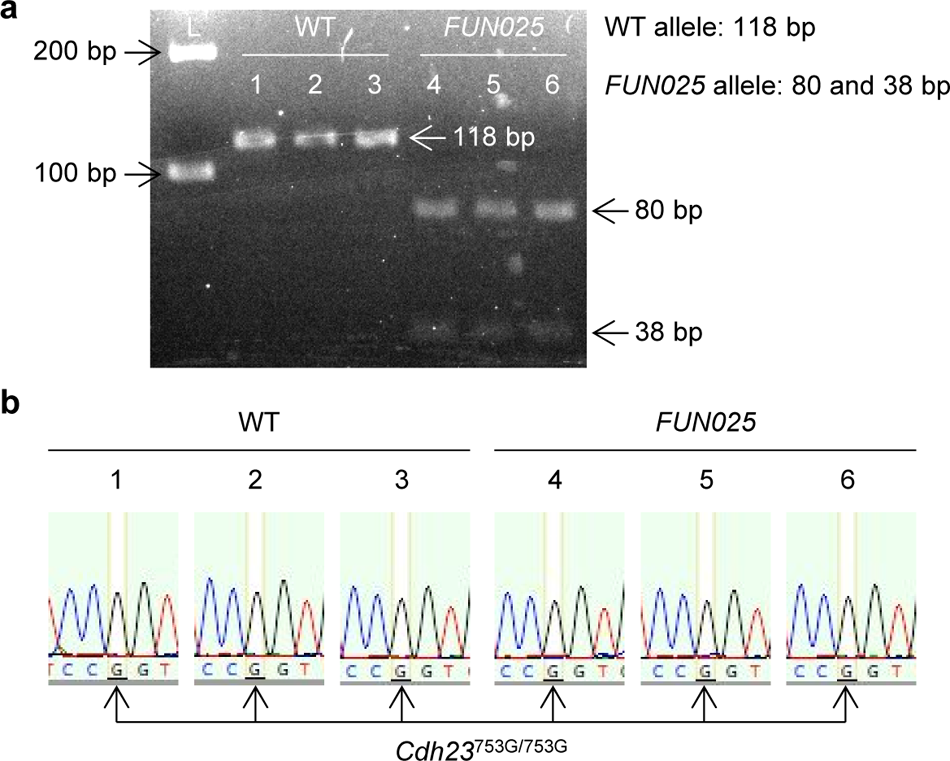
Genotyping of WT and *FUN025* mutant mice. ***a***, *FUN025* genotyping: PCR primers were used for amplification of the WT allele (118 bp) and the *FUN025* allele (118 bp). Subsequently, Cac8I was used to digest the *FUN025* allele specifically to generate two bands (80 and 38 bp). L, 100 bp DNA ladder. ***b***, *Cdh23* genotyping: The region of DNA containing the 753^rd^ nucleotide in *Cdh23* was sequenced in WT and *FUN025* mice. All the mice examined had the same wild-type *Cdh23* genotype (*Cdh23*^753G/753G^). *Cdh23*, cadherin 23. Data are shown as means ± SEM. WT, *Tmem135*^+/+^; *FUN025*, *Tmem135*^FUN025/FUN025^.

#### Cdh23 genotyping

The *Tmem135*^FUN025/+^ mice were developed on the C57BL/6J background (Lee et al., 2016). The C57BL/6J mouse strain carries a single-nucleotide polymorphism (G→A) at nucleotide 753 in exon 7 of *Cdh23* (cadherin 23) that causes early-onset age-related hearing loss (ARHL) by 9-12 months of age (Noben-Trauth et al., 2003; Han and Someya, 2013). To replace this ARHL-susceptibility allele (*Cdh23*^753A^) with the AHL-resistance allele (*Cdh23*^753G^), the C57BL/6J-*Tmem135*^FUN025/+^ mice were backcrossed for four generations onto the CBA/CaJ mouse strain, a normal-hearing strain that does not carry the G→A polymorphism (Noben-Trauth et al., 2003). Genomic DNA was isolated from the tails of CBA/CaJ-*Tmem135*^+/+^ and CBA/CaJ-*Tmem135*^FUN025/FUN025^ mice. Primer sequences for PCR were as follows (Kim et al., 2019; Kim et al., 2020):

Forward 5’-GATCAAGACAAGACCAGACCTCTGTC-3’;

Reverse 5’-GAGCTACCAGGAACAGCTTGGGCCTG-3’.

These primers were used for amplification of the AHL-resistance allele (*Cdh23*^753G^) (360 bp) and the ARHL-susceptibility allele (*Cdh23*^753A^) (360 bp). Subsequently, the region of DNA containing the 753^rd^ nucleotide in *Cdh23* were Sanger sequenced in WT and *FUN025* mice (Fig. 1*b*). All the mice examined had the same wild-type *Cdh23* genotype (*Cdh23*^753G/753G^) (Fig. 1*b*). Cycling conditions for PCR were as follows: 95 °C for 2 min; 35 cycles of 95 °C for 30 sec, 60 °C for 1 min, 72 °C for 1 min; 72 °C for 5 min.

### 2.3. ABR

Auditory brainstem response (ABR) tests were performed using the TDT system with RZ6 hardware and BioSigRZ software (Tucker-Davis Technologies) according to the manufacturer’s ABR User Guide and as previously described (Kim et al., 2019; Kim et al., 2020; Kim et al., 2023). Mice were anesthetized with ketamine (100 mg/kg) and xylazine (10 mg/kg) by intraperitoneal injection. The needle electrodes connected to the RA4PA-RA4LI (Tucker-Davis Technologies) were subdermally inserted at the vertex (channel 1), ipsilateral ear (right ear, reference), and contralateral ear (left ear, ground). The tube connected to one MF1 speaker (Tucker-Davis Technologies) was positioned in the right ear canal. ABR waveforms were collected in response to tone sound stimuli at 8, 16, 32, 48, and 64 kHz. At each frequency, the sound level was reduced in 5-10 dB SPL steps from 90 to 10 dB SPL. The ABR threshold was defined as the lowest stimulus level at which the ABR wave I was reproducibly identified. Four age groups, two sex groups, and three genotype groups were used in the ABR tests: 6-month-old female (10 total, 6 Het and 4 *FUN025*), 1-month-old male (12 total, 6 WT and 6 *FUN025*), 1-month-old female (10 total, 5 WT and 5 *FUN025*), 3-month-old male (15 total, 7 WT and 8 *FUN025*), 3-month-old female (12 total, 7 WT and 5 *FUN025*), 12-month-old male (17 total, 7 WT and 10 *FUN025*), and 12-month-old female (16 total, 8 WT and 8 *FUN025*). The same animals were used for the DPOAE and rotarod performance experiments.

### 2.4. DPOAE

Distortion product otoacoustic emission (DPOAE) tests were performed using the TDT system with RZ6 hardware and BioSigRZ software (Tucker-Davis Technologies) according to the manufacturer’s DPOAE User Guide. Mice were anesthetized with ketamine (100 mg/kg) and xylazine (10 mg/kg) by intraperitoneal injection. The tip connected to the ER10B+ (Etymotic Research) and two MF1 speakers (Tucker-Davis Technologies) was positioned in the right ear canal. DPOAE amplitudes at 2F1-F2 were collected in response to two-tone sound stimuli. The tone frequencies F1 and F2 had a F2/F1 ratio of 1.2 and were geometrically centered about 8, 16, and 32 kHz. At each center frequency, the tone levels L1 and L2 remained equal and were reduced in 10 dB steps from 80 to 20 dB SPL. The DPOAE threshold was defined as the lowest stimulus level at which the 2F1-F2 distortion product was more than 6 dB SPL above the noise floor (Powers et al., 2006). The noise floor was defined as the average of the distortion products at ten neighboring frequencies (five above and five below the 2F1-F2) (Powers et al., 2006). One age group, two sex groups, and two genotype groups were used in the DPOAE tests: 4-month-old male (14 total, 8 WT and 6 *FUN025*) and 4-month-old female (16 total, 8 WT and 8 *FUN025*).

### 2.5. Rotarod latency

Mice were trained to use a rotarod apparatus (Rota Rod Rotamex 5, Columbus Instruments) with an acceleration of 2 rpm every 17 sec from 4 rpm to 40 rpm during the first five trials (trials 1-5). The rotarod performance (average latency to fall) was analyzed during the next three trials (trials 6-8) as previously described (Kim et al., 2023). One age group, two sex groups, and two genotype groups were used in the rotarod latency measurements; 3-month-old male (19 total, 9 WT and 10 *FUN025*) and 3-month-old female (17 total, 7 WT and 10 *FUN025*).

### 2.6. Cochlear histopathology

#### Sample preparation

Mice were sacrificed and temporal bones were extracted from the head. For hair cell (HC) survival assessment, cochleae were fixed in 4% paraformaldehyde (PFA), decalcified in 10% ethylenediaminetetraacetic acid (EDTA), and microdissected as previously described (Honkura et al., 2019; Hemmi et al., 2023). For spiral ganglion neuron (SGN) density and stria vascularis (SV) thickness measurements, cochleae were fixed in 4% PFA, decalcified in 10% EDTA, embedded in paraffin, cut at 3 μm, mounted on glass slides, and stained with hematoxylin and eosin (H&E) as previously described (Honkura et al., 2019; Hemmi et al., 2023).

#### Hair cell survival

Inner hair cells (IHCs) and outer hair cells (OHCs) were counted as previously described (Honkura et al., 2019; Hemmi et al., 2023). The hair cells were stained for F-actin with rhodamine-conjugated phalloidin (1:100, Rhodamine Phalloidin, Thermo Fisher Scientific) for 1 h at room temperature, and mounted on glass slides with VECTASHIELD Antifade Mounting Medium with DAPI (Vector Laboratories, H-1200-10). Low-power fluorescence images were obtained using a microscope (BZ-9000, Keyence, Osaka, Japan) and BZ-H1 software (Keyence), and the full-length cochlea image was assembled and analyzed in Adobe Photoshop. An ImageJ plug-in (National Institutes of Health) plug-in was used to create a cochlear frequency map (https://meeeplfiles.partners.org/Measure_line.class). Damaged and undamaged IHCs and OHCs were counted at 8, 11.3, 16, 22.6, 32, and 45.2 kHz cochlear frequency regions. Two age groups, one sex group, and two genotype groups were used in the hair cell counting: 2-month-old male (10 total, 5 WT and 5 *FUN025*) and 13-month-old male (8 total, 3 WT and 5 *FUN025*).

#### SGN density

Spiral ganglion neurons (SGNs) were counted as previously described (Honkura et al., 2019; Hemmi et al., 2023). H&E-stained cochlear sections were imaged using a microscope (BZ-9000, Keyence) with a 40x objective. The number of SGNs and the area were measured in the apical, middle, and basal cochlear regions using ImageJ. Three sections of the apical, middle, and basal cochlear regions were examined per mouse. Two age groups, one sex group, and two genotype groups were used in the SGN density measurements: 2-month-old male (10 total, 5 WT and 5 *FUN025*) and 13-month-old male (8 total, 3 WT and 5 *FUN025*).

#### SV thickness

Stria vascularis (SV) thickness was measured as previously described (Honkura et al., 2019; Hemmi et al., 2023). H&E-stained cochlear sections were imaged using a microscope (BZ-9000, Keyence) with a 40x objective. The thickness of SV was measured in the apical, middle, and basal cochlear regions using the ImageJ. The measurement was made by using a cursor to draw a line from the marginal cells to the basal cells halfway between the Reissner’s membrane and the spiral prominence. Three sections of the apical, middle, and basal cochlear regions were examined per mouse. Two age groups, one sex group, and two genotype groups were used in the SV thickness measurements: 2-month-old male (10 total, 5 WT and 5 *FUN025*) and 13-month-old male (8 total, 3 WT and 5 *FUN025*).

### 2.7. Immunofluorescence

Mice were sacrificed and temporal bones were extracted from the head. Cochleae were fixed in 4% PFA in phosphate-buffered saline (PBS) for 2 h at room temperature, decalcified in 0.12 M EDTA in PBS for 2 days at room temperature, and microdissected as previously described (Suzuki et al., 2016; Suzuki et al., 2017). Microdissected cochlear pieces were blocked with 20% normal goat serum, 0.3% Triton X-100 in PBS for 1 h at room temperature followed by overnight incubation at 37 °C with mouse anti-MYO7A (myosin VIIA) (1:200, Santa Cruz Biotechnology, sc-74516). Cochlear pieces were then incubated with fluorophore-conjugated species-appropriate secondary antibodies (Thermo Fisher Scientific) for 1 hour at 37 °C for 15 min at room temperature, and mounted on glass slides with ProLong Glass Antifade Mountant (Thermo Fisher Scientific). Immunostained cochlear whole mounts were visualized using an inverted microscope (Nikon Ti2) with either 60x (CFI Plan Apo, 1.4 N.A.) or 100x (CFI Apo TIRF 1.49 N.A.) oil immersion objectives and imaged using a spinning disk confocal scanner (Yokogawa CSU-X1) with Live-SR (Gataca Systems) and Prime 95B sCMOS camera (Teledyne Photometrics). Image stacks were captured with a 0.2 μm spacing. Image stacks were captured with 0.2 μm spacing and analyzed using NIS-Elements (Nikon) and ImageJ.

### 2.8. RNA in situ hybridization

RNA *in situ* hybridization was conducted using the BaseScope assay (Advanced Cell Diagnostics) according to the manufacturer’s instructions with a modified protocol previously described (Ghosh et al., 2022). Mouse cochleae were fixed in 4% PFA for 3 hours and microdissected in cold 20% RNAlater (Thermo Fisher Scientific) in PBS. Microdissected samples were stored in RNAlater at 4 °C. After removing RNAlater, samples were incubated with 50% ethanol for 5 min at room temperature followed by incubation with 70%, 95%, and 100% ethanol. After removing 100% ethanol, samples were incubated with RNAscope Hydrogen Peroxide (Advanced Cell Diagnostics) for 30 min at room temperature and washed with Milli-Q water. Samples were then incubated with Antigen Unmasking Solution, Citrate-Based (1:100, Vector Laboratories) for 5 min at 65 °C and washed. Samples were then incubated with RNAscope Protease Plus (Advanced Cell Diagnostics) for 30 min at room temperature, washed, and stored in Milli-Q water at 4 °C. After removing Milli-Q water, samples were incubated with BaseScope probes for 2 hours at 40 °C. The following BaseScope probes were used: BA-Mm-Tmem135-E11E12-C1 (BaseScope Target Probe targeting 1191-1235 of NM_028343.5, 1256981-C1); BA-Mm-Ppib-1zz (Positive Control Probe, 712351); BA-DapB-1zz (Negative Control Probe, 701021). After washing with RNAscope Wash Buffer (1:50, Advanced Cell Diagnostics), samples were incubated with BaseScope v2 AMP1-8 (Advanced Cell Diagnostics) followed by washing between each amplification step. After washing with RNAscope Wash Buffer, samples were incubated with a mixture of BaseScope Fast RED-A (Advanced Cell Diagnostics) and BaseScope Fast RED-B (Advanced Cell Diagnostics) at a 60:1 ratio for 10 min at room temperature and washed with Milli-Q water. To conduct co-staining for hair cells and nuclei, samples were incubated with rabbit anti-MYO7A (Proteus BioSciences, 25-6790) in 0.1% Triton X-100 overnight at 4 °C. Samples were then incubated with donkey anti-rabbit conjugated with Alexa Fluor 488 in 0.1% Triton X-100 for 2 hours at room temperature, stained with DAPI (1.5 μg/mL) for 15 min at room temperature, mounted on glass slides with ProLong Glass Antifade Mountant (Thermo Fisher Scientific), and imaged using a spinning disk confocal microscope as described above.

### 2.9. Statistics

Two-way ANOVA (analysis of variance) with Bonferroni’s multiple comparisons test was carried out using Prism v9.0 (GraphPad Software) to analyze ABR thresholds, DPOAE thresholds, DPOAE amplitudes, IHC survivals, OHC survivals, SGN densities, and SV thicknesses. Unpaired two-tailed Student’s *t*-test were performed using Prism to analyze rotarod latencies.

## 3. Results

### 3.1. *FUN025* mutant mice display progressive hearing loss

*FUN025* mice were isolated through fundus examination in an ENU (N-ethyl-N-nitrosourea) mouse mutagenesis project (Pinto et al., 2004). A subsequent study by Ikeda and co-workers revealed a point mutation (T>C) in the splice-donor site adjacent to exon 12 of *Tmem135* in *FUN025* mice (Lee et al., 2016). This mutation disrupts the splice donor site, resulting in skipping of exon 12 and a frame shift creating a premature stop codon in *FUN025* mice. A knock-out allele of *Tmem135* exhibits the same retinal phenotypes as the *FUN025* mutant mice, indicating that the *FUN025* mutation is likely a loss-of-function allele (Lee et al., 2016).

Because *FUN025* mutant mice were originally generated in the C57BL/6J background (Lee et al., 2016), they are homozygous for the ARHL-susceptibility allele (*Cdh23*^753A^) (Noben-Trauth et al., 2003). To avoid this confounding variable in our experiments, we crossed the *FUN025* mutant mouse for 4 generations onto the CBA/CaJ mouse strain, a normal-hearing strain homozygous for the ARHL-resistance allele (*Cdh23*^753G^) (Kim et al., 2019). We genotyped N4 CBA/CaJ-*Tmem135*^+/+^ (WT) and CBA/CaJ-*Tmem135*^FUN025/FUN025^ (*FUN025*) mice for *FUN025* (Fig. 1*a*) and *Cdh23* (Fig. 1*b*). We confirmed that all WT and *FUN025* mice examined had the same wild-type *Cdh23* genotype (*Cdh23*^753G/753G^) (Fig. 1*b*).

To evaluate auditory function, we measured ABR thresholds at 8, 16, 32, 48, and 64 kHz in female heterozygous *FUN025* and homozygous *FUN025* mice at 6 months of age (Fig. 2*a*). Homozygous *FUN025* mice displayed a 15-33 dB SPL increase in ABR thresholds at 8, 16, 48, and 64 kHz compared to heterozygous *FUN025* littermates (Fig. 2a). Because heterozygous *FUN025* mice did not display elevated ABR thresholds at 8, 16, 32, and 48 kHz, subsequent studies were conducted in WT and *FUN025* mice. We measured ABR thresholds at 8, 16, 32, 48, and 64 kHz in male and female WT and *FUN025* mice at 1, 3, and 12 months of age (Fig. 2*b*-*g*). At 1 month of age, there were no differences in ABR thresholds at all the frequencies measured between WT and *FUN025* mice in males (Fig. 2*b*) or females (Fig. 2*e*), indicating normal hearing. However, at 3 months of age, male *FUN025* mice displayed a 22-38 dB SPL increase in ABR thresholds at 8, 16, 48, and 64 kHz (Fig. 2*c*). Female *FUN025* mice also displayed a 23-40 dB SPL increase in ABR thresholds at 8, 16, and 64 kHz at 3 months of age (Fig. 2*f*). At 12 months of age, both male and female *FUN025* mice displayed profound hearing loss (Fig. 2*d*, *g*). There were no observed sex differences in ABR thresholds at all the frequencies measured in WT or *FUN025* mice at 1, 3, or 12 months of age.

**Figure 2.**
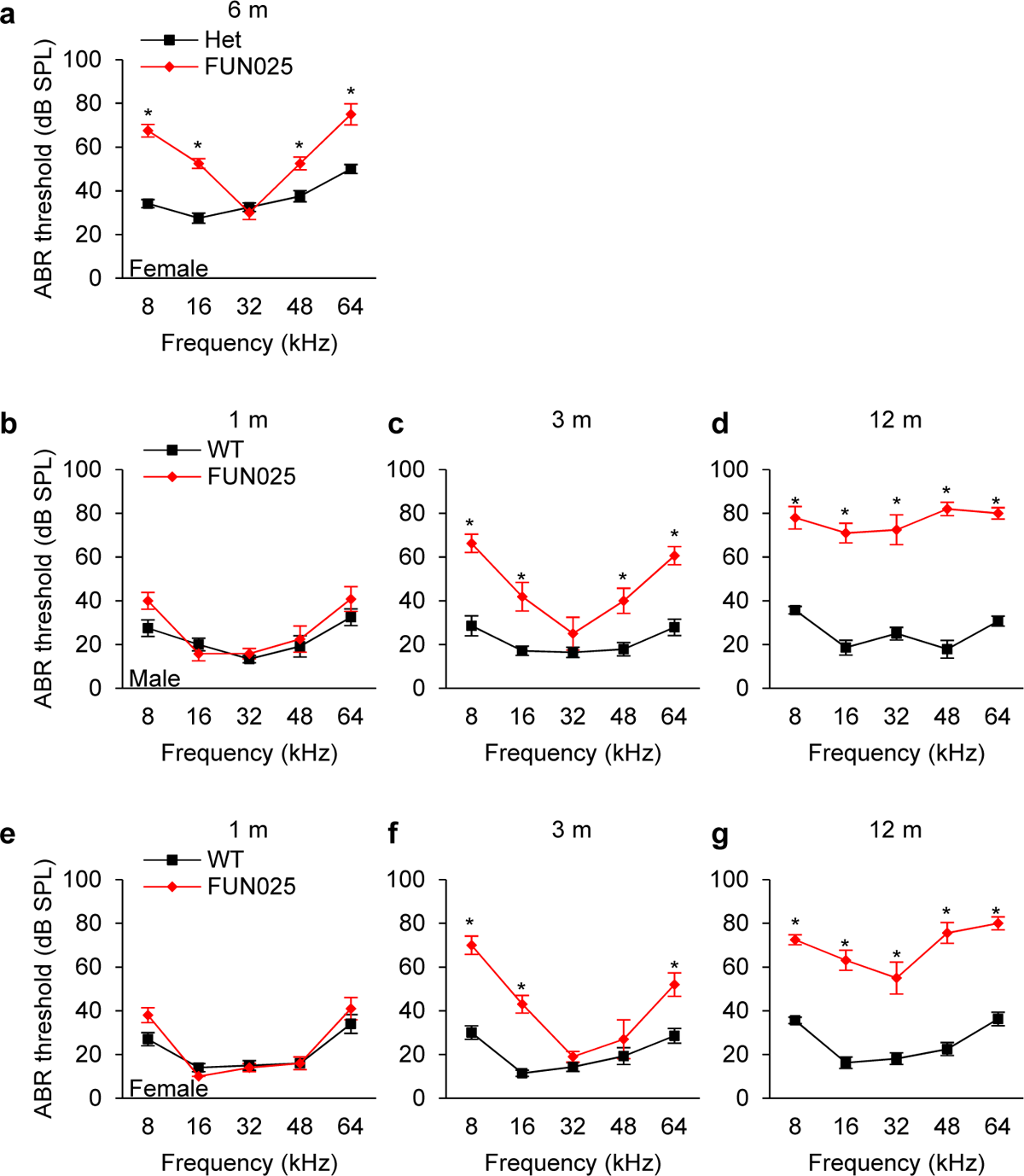
*FUN025* mutant mice display progressive profound hearing loss. ***a***, ABR thresholds were measured at 8, 16, 32, 48, and 64 kHz in female Het and *FUN025* mice at 6 months of age; 6-month-old female (10, 6 Het and 4 *FUN025*). ***b-g***, ABR thresholds were measured at 8, 16, 32, 48, and 64 kHz in male (***b-d***) and female (***e-g***) WT and *FUN025* mice at 1, 3, and 12 months of age; 1-month-old male (12, 6 WT and 6 *FUN025*), 1-month-old female (10, 5 WT and 5 *FUN025*), 3-month-old male (15, 7 WT and 8 *FUN025*), 3-month-old female (12, 7 WT and 5 *FUN025*), 12-month-old male (17, 7 WT and 10 *FUN025*), and 12-month-old female (16, 8 WT and 8 *FUN025*). Data are shown as means ± SEM. Two-way ANOVA with Bonferroni’s multiple comparisons tests were performed. *, *P* < 0.05, Het vs. *FUN025*, WT vs. *FUN025*. m, month; WT, *Tmem135*^+/+^; Het, *Tmem135*^FUN025/+^; *FUN025*, *Tmem135*^FUN025/FUN025^.

To evaluate auditory dysfunction in more detail, we measured DPOAE thresholds and amplitudes at 8, 16, and 32 kHz in male and female WT and *FUN025* mice at 4 months of age (Fig. 3*a-h*). Male *FUN025* mice displayed a 20-30 dB SPL increase in DPOAE thresholds at 8 and 16 kHz compared to WT mice (Fig. 3*a*). Similarly, female *FUN025* mice displayed a 25-26 dB SPL increase in DPOAE thresholds at 8 and 16 kHz compared to WT mice (Fig. 3*e*). Both male and female *FUN025* mice displayed significantly smaller DPOAE amplitudes at 8 and 16 kHz compared to WT mice (Fig. 3*b-c*, *f-g*). There were no observed sex differences in DPOAE thresholds or amplitudes at all the frequencies and sound levels measured in WT or *FUN025* mice.

**Figure 3.**
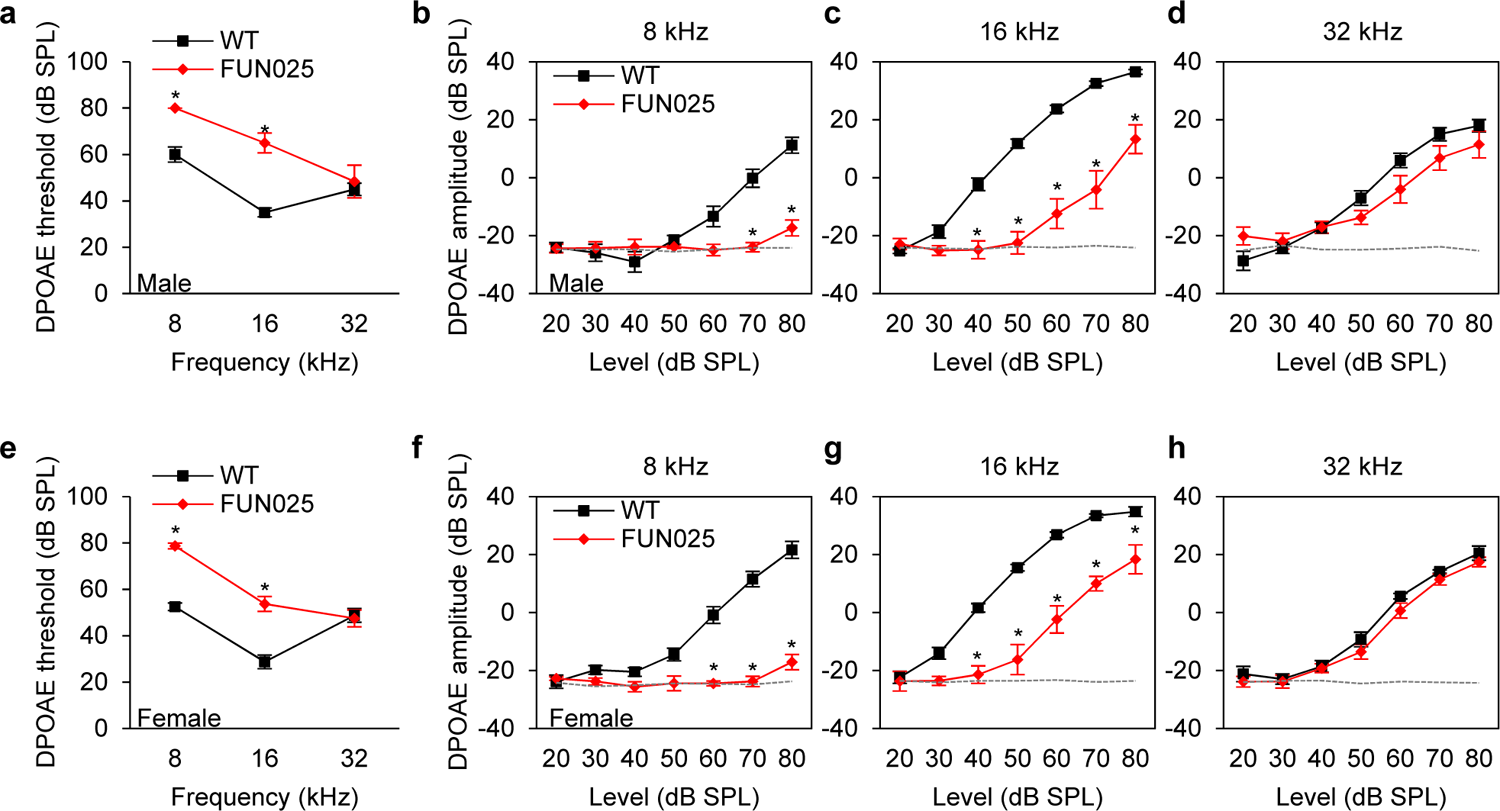
Young *FUN025* mutant mice display increased DPOAE thresholds and decreased DPOAE amplitudes. ***a-h***, DPOAE thresholds and amplitudes were measured at 8, 16, and 32 kHz in male (***a-d***) and female (***e-h***) WT and *FUN025* mice at 4 months of age; 4-month-old male (14, 8 WT and 6 *FUN025*) and 4-month-old female (16, 8 WT and 8 *FUN025*). Data are shown as means ± SEM. Two-way ANOVA with Bonferroni’s multiple comparisons tests were performed. *, *P* < 0.05, WT vs. *FUN025*. Dashed lines present noise floors. WT, *Tmem135*^+/+^; *FUN025*, *Tmem135FUN025/FUN025*.

To assess vestibular function in *FUN025* mutant mice, we performed rotarod balance tests on male and female WT and *FUN025* mice at 3 months of age (Fig. 4*a-b*). There were no differences in rotarod latencies between WT and *FUN025* mice in males (Fig. 4*a*) or females (Fig. 4*b*), indicating that the *FUN025* mutation in *Tmem135* does not grossly affect vestibular function. Together, these physiological test results demonstrate that TMEM135 is necessary for maintaining normal auditory function.

**Figure 4.**
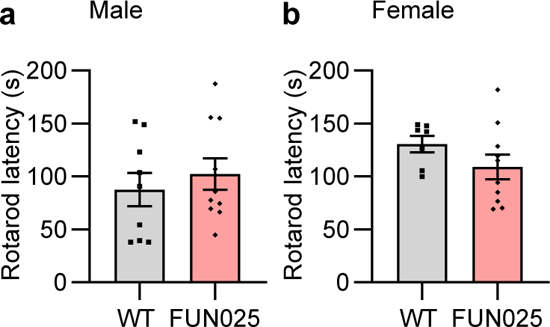
The *FUN025* mutation in *Tmem135* does not affect rotarod balance performance in young mice. ***a-b***, Rotarod latencies were measured in male (***a***) and female (***b***) WT and *FUN025* mice at 3 months of age; 3-month-old male (19, 9 WT and 10 *FUN025*) and 3-month-old female (17, 7 WT and 10 *FUN025*). Data are shown as means ± SEM. Unpaired two-tailed Student’s *t*-test were performed. WT, *Tmem135*^+/+^; *FUN025*, *Tmem135*^FUN025/FUN025^.

### 3.2. Tmem135 is expressed in the inner and outer hair cells

At 3-4 months of age, *FUN025* mice displayed a 20-40 dB SPL increase in ABR thresholds and a 20-30 dB increase in DPOAE thresholds compared to WT mice. At 12 months of age, *FUN025* mice displayed profound hearing loss; we therefore hypothesized *Tmem135* was expressed in hair cells. To test this, we performed BaseScope *in situ* hybridization to detect *Tmem135* transcripts in cochlear whole mount tissues from P90 C57BL/6J wild-type mice (Fig. 5*a-c*). The signals from the *Tmem135* BaseScope probe were detected in IHCs, OHCs, and supporting cells (Fig. 5*a*). These results suggest that TMEM135 is necessary for maintaining hair cells in the aging cochlea.

**Figure 5.**
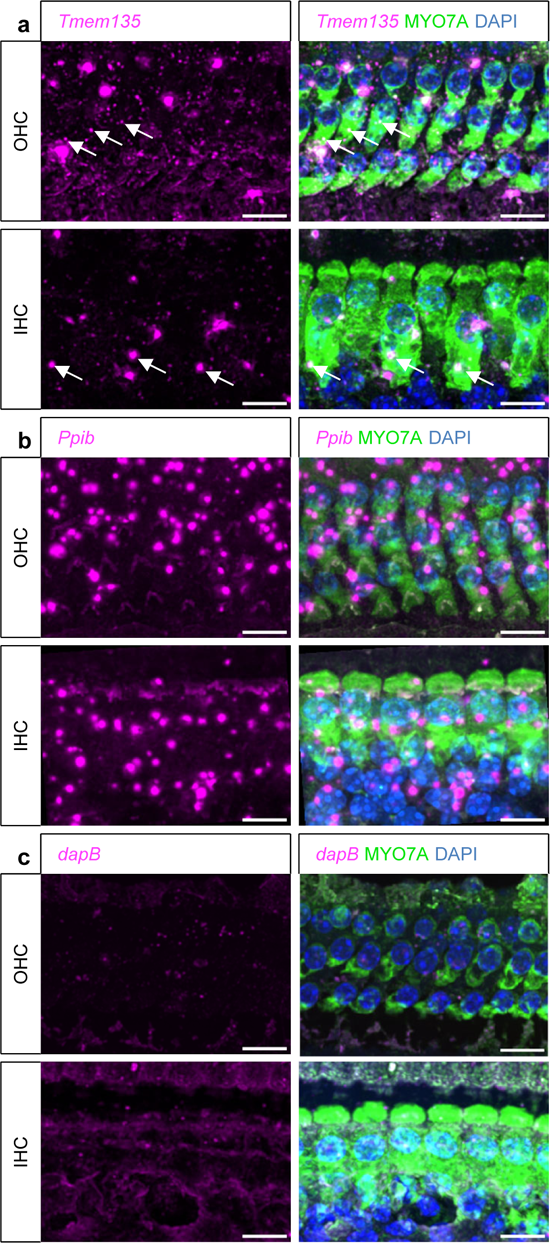
*Tmem135* mRNA is expressed in cochlear hair cells. ***a***, Maximal projection confocal images illustrating the BaseScope signals for *Tmem135* in the cochlear whole mount tissue from a 3-month-old C57BL/6J mouse. The localization of the *Tmem135* BaseScope signals (magenta) in cochlear hair cells is demonstrated through co-staining for hair cells (anti-MYO7A, green) and nuclei (DAPI, blue). ***b***, Positive control with the mouse *Ppib* probe. *Ppib*, peptidylprolyl isomerase B. ***c***, Negative control with the bacterial *dapB* probe. *dapB*, 4-hydroxy-tetrahydrodipicolinate reductase. Scale bar = 10 μm. MYO7A, myosin VIIA; DAPI, 4′,6-diamidino-2-phenylindole; IHC, inner hair cell; OHC, outer hair cell.

### 3.3. FUN025 mutant mice display progressive loss of hair cells and spiral ganglion neurons in the cochlea

The major sites of cochlear pathology generally include HCs, SGNs, and/or SV (Yamasoba et al., 2013). To explore why loss of *Tmem135* resulted in hearing loss, we quantified IHC and OHC numbers at 8, 11.3, 16, 22.6, 32, and 45.2 kHz regions in the cochlear whole mount tissues from male WT and *FUN025* mice at 2 and 13 months of age (Fig. 6*a-n*). Because there were no sex differences in ABRs or DPOAEs in WT or *FUN025* mutant mice, subsequent cochlear tissue studies were conducted in either males or females. At 2 months of age, while there were no differences in IHC survival at all the cochlear frequency regions measured between WT and *FUN025* mice (Fig. 6*a-f*, *m*, Fig. *7a-b, e-f*), *FUN025* mice displayed a 46-66% decrease in OHC survival at 8, 11.3, 16, and 22.6 kHz cochlear regions compared to WT mice (Fig. 6*g-j*, *n*, Fig. *7a-b*, *e-f*). At 13 months of age, *FUN025* mice displayed a 23-27% decrease in IHC survival at 16, 22.6, and 45.2 kHz cochlear regions (Fig. 6*c*, *d*, *f*, *m*, Fig. 7*c-d*) and near-total loss of OHCs at 8, 11.3, 16, 22.6, and 32 kHz cochlear regions compared to WT mice (Fig. 6*g-k*, *n*, Fig. 7*c-d*), indicating that the *FUN025* mutation causes the progressive loss of both OHCs and IHCs.

**Figure 6.**
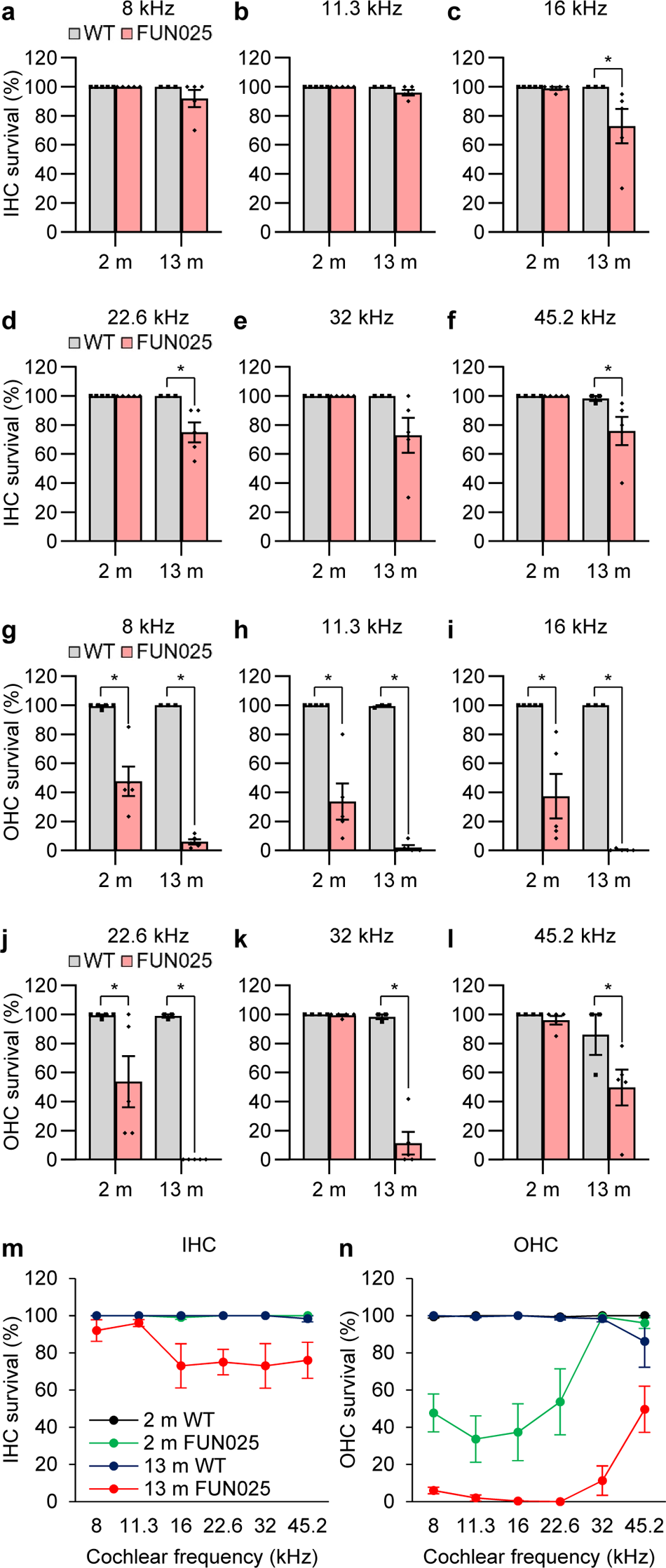
*FUN025* mutant mice display progressive profound loss of cochlear hair cells. ***a-n***, Inner (***a-f***, ***m***) and outer (***g-l***, ***n***) hair cell numbers were quantified at 8, 11.3, 16, 22.6, 32, and 45.2 kHz regions in the cochlear whole mount tissues from male WT and *FUN025* mice at 2 and 13 months of age; 2-month-old male (10, 5 WT and 5 *FUN025*) and 13-month-old male (8, 3 WT and 5 *FUN025*). Data are shown as means ± SEM. Two-way ANOVA with Bonferroni’s multiple comparisons tests were performed. *, *P* < 0.05, WT vs. *FUN025*. IHC, inner hair cell; OHC, outer hair cell; m, month; WT, *Tmem135*^+/+^; *FUN025*, *Tmem135*^FUN025/FUN025^.

**Figure 7.**
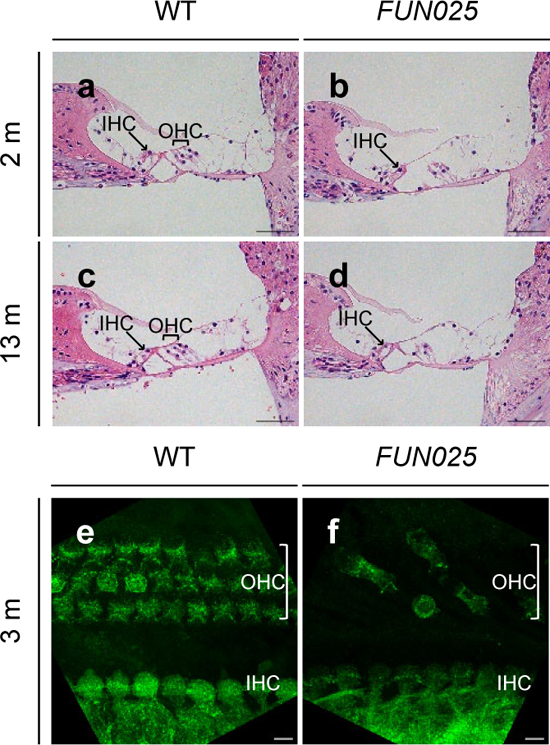
Representative images of cochlear hair cells in WT and *FUN025* mutant mice. ***a-d***, Images were obtained in the middle region of the cochleae from male WT and *FUN025* mice at 2 and 13 months of age. Paraffin sections of cochlear tissues were stained with H&E. 40x images were obtained. Scale bar = 50 μm. H&E, hematoxylin, and eosin**. *e-f***, Images were obtained in the middle region of the cochleae from 3-month-old male WT (***e***) and *FUN025* (***f***) mice. Cochlear whole mount tissues were stained for hair cells (anti-MYO7A, green). Maximum intensity projections of confocal z-stacks (0.2 μm step size, 60x objective) were obtained. Scale bar = 5 μm. MYO7A, myosin VIIA; IHC, inner hair cell; OHC, outer hair cell; m, month; WT, *Tmem135*^+/+^; *FUN025*, *Tmem135*^FUN025/FUN025^.

To further characterize how the *FUN025* mutation causes hearing loss, we quantified SGN densities in the apical, middle, and basal regions of the cochlear tissue sections from male WT and *FUN025* mice at 2 and 13 months of age (Fig. 8*a-c*). At 2 months of age, *FUN025* mice displayed a 19% decrease in SGN densities in the middle cochlear region compared to WT mice (Fig. 8*b*). At 13 months of age, *FUN025* mice displayed a 29-48% decrease in SGN densities in the apical, middle, and basal cochlear regions compared to WT mice (Fig. 8*a-c*), indicating that the *FUN025* mutation causes progressive degeneration of SGNs, consistent with the HC survival results.

**Figure 8.**
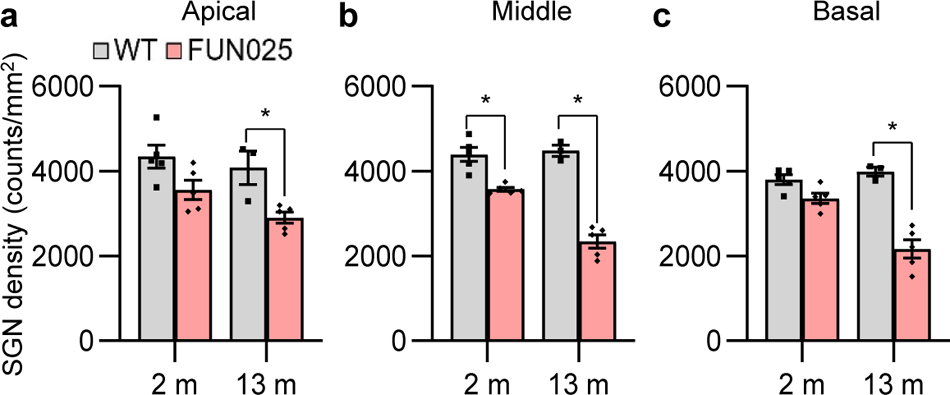
*FUN025* mutant mice display progressive profound loss of spiral ganglion neurons. ***a-c***, Spiral ganglion neuron densities were measured in the apical, middle, and basal regions in the cochlear tissue sections from male WT and *FUN025* mice at 2 and 13 months of age; 2-month-old male (10, 5 WT and 5 *FUN025*) and 13-month-old male (8, 3 WT and 5 *FUN025*). Data are shown as means ± SEM. Two-way ANOVA with Bonferroni’s multiple comparisons tests were performed. *, *P* < 0.05, WT vs. *FUN025*. SGN, spiral ganglion neuron; m, month; WT, *Tmem135*^+/+^; *FUN025*, *Tmem135FUN025/FUN025*.

Finally, we measured SV thicknesses in the apical, middle, and basal regions of the cochlear tissue sections from male WT and *FUN025* mice at 2 and 13 months of age (Fig. 9*a-c*). At 2 months of age, *FUN025* mice displayed a 20-24% decrease in SV thicknesses in the apical and middle cochlear regions compared to WT mice (Fig. 9*a-b*). At 13 months of age, *FUN025* mice displayed a 29-54% decrease in SV thicknesses in the apical, middle, and basal cochlear regions compared to WT mice (Fig. 9*a-c*), indicating that the *FUN025* mutation causes SV atrophy. Together, these histological findings demonstrate that the *FUN025* mutation in *Tmem135* causes a range of pathological changes that contribute to hearing loss, including the near-total loss of OHCs, and more modest changes in SGN density and SV integrity.

**Figure 9.**
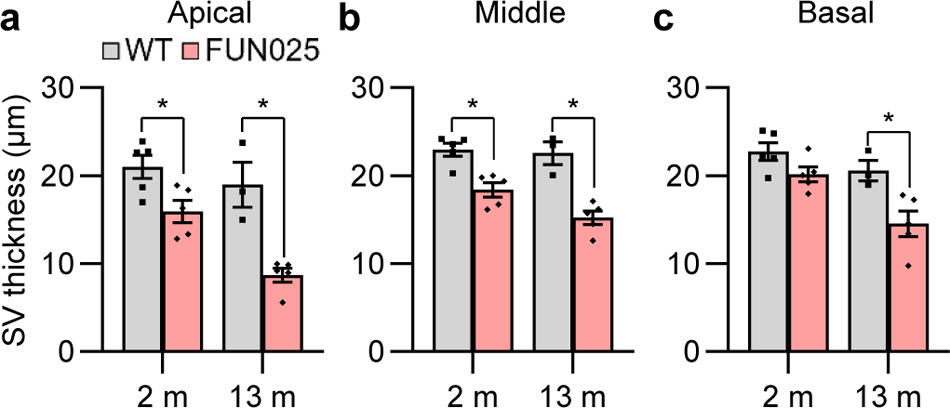
*FUN025* mutant mice display progressive profound loss of stria vascularis cells. ***a-c***, Stria vascularis thicknesses were measured in the apical, middle, and basal regions in the cochlear tissue sections from male WT and *FUN025* mice at 2 and 13 months of age; 2-month-old male (10, 5 WT and 5 *FUN025*) and 13-month-old male (8, 3 WT and 5 *FUN025*). Data are shown as means ± SEM. Two-way ANOVA with Bonferroni’s multiple comparisons tests were performed. *, *P* < 0.05, WT vs. *FUN025*. SV, stria vascularis; m, month; WT, *Tmem135*^+/+^; *FUN025*, *Tmem135FUN025/FUN025*.

## 4. Discussion

Mutations in *TMEM135* have not previously been associated with inherited forms of human deafness. In this report, we have demonstrated that a loss of function allele in *Tmem135* (*FUN025*) causes a progressive sensorineural hearing loss in CBA/CaJ mice. At one month of age, there were no differences in ABR thresholds at 8, 16, 32, 48, or 64 kHz between WT and *FUN025* males or WT and *FUN025* females, indicating normal hearing. However, by 3 months of age, both male and female *FUN025* mutant mice displayed significantly elevated ABR thresholds at 8, 16, and 64 kHz. By 12 months of age, both male and female *FUN025* mice displayed profound hearing loss. Middle-aged *FUN025* mice also exhibited near-total loss of OHCs, severe to profound degeneration of SGNs, and severe SV atrophy in the apical, middle, and basal cochlear regions compared to WT mice, validating the physiological test results. Additionally, using the same *FUN025* mutant mice, a previous study has also demonstrated that the *FUN025* mutation in *Tmem135* causes age-related retinal pathologies (Lee et al., 2016). Together, these studies and our findings suggest that loss of *TMEM135* accelerates the progression of age-related hearing and vision impairments.

The broad range of cochlear pathology observed in *Tmem135* mutant mice suggests that TMEM135 has key roles in multiple cell types. Consistent with this, our BaseScope results show expression in OHCs, IHCs as well as supporting cells. These results build upon prior transcriptomic analyses that detected *Tmem135* in IHCs, OHCs, pillar cells, Deiters’ cells, and/or stria vascularis in CBA/J mice at approximately P30 (Liu et al., 2018; Korrapati et al., 2019) and CD1 mice at P1 (Kolla et all., 2020). Our results show that hair cells expressing *Tmem135* are lost in the aging *FUN025* mouse. Hair cells are well documented to have elevated energetic demand and thought to have a high concentration of mitochondria within their cytoplasm (Fettiplace R et al., 2019; Lysakowski et al., 2022). Given the functions of TMEM135 in regulating mitochondrial dynamics (Lee et al., 2016) and lipid metabolism in other cell types (Landowski et al., 2022), we speculate that TMEM135 is similarly critical for mitochondrial function in these highly metabolically-active cochlear cells. Whilst we cannot exclude an indirect effect of stria pathology and potential disruption to the endocochlear potential may contribute to the loss of OHCs in *FUN025* mice (Liu et al., 2016), future studies will address the localization of TMEM135 protein in individual cochlear cell types and study the effects of *Tmem135* deletion in specific cell types.

In summary, we demonstrate a progressive hearing loss phenotype caused by a mutation in *Tmem135*. In addition to identifying a new protein required for the maintenance of key cochlear cell types during normal aging of the mouse cochlea, our work highlights *TMEM135* as a potential candidate gene in studies of hereditary hearing loss and age-related hearing impairment (ARHI).

## Acknowledgments

This research was supported by R01 DC012552 (Someya), R01 DC014437 (Someya) and R01 DC018827 (Bird) from the National Institutes of Health / National Institute on Deafness and Communication Disorders, the Claude D. Pepper Older Americans Independence Centers at the University of Florida (P30 AG028740) from the National Institutes of Health / National Institute on Aging and Evelyn F. McKnight Brain Research Foundation.

## Data availability

Data will be made available on request.

## Author contributions

**Mi-Jung Kim**: Writing – original draft, Writing – review & editing, performed physiological (ABR and DPOAE) and histological experiments, analyzed and collected data, **Shion Simms**: performed genotyping and physiological experiments, analyzed and collected data, Writing – review & editing, **Ghazaleh Behnammanesh**: performed BaseScope experiments, analyzed and collected data, Writing – review & editing, **Yohei Honkura**: performed histological experiments (SGN and SV thickness measurements), Writing – review & editing, **Jun Suzuki**: performed histological experiments (hair cell counts), Writing – review & editing, **Hyo-Jin Park**: performed physiological experiments (ABR), **Marcus Milani**: performed physiological experiments (rotarod test), **Yukio Katori**: Writing – review & editing, Supervision, Funding acquisition, Writing – review & editing, **Jonathan E Bird**: performed BaseScope experiments, Writing – review & editing, Supervision, Funding acquisition, **Akihiro Ikeda**: performed backcrossing, provide the *FUN025* mice, Supervision, and Writing – review & editing, **Shinichi Someya**: Writing – original draft, Writing – review & editing, Supervision, Project administration, Funding acquisition.

